# Whole genome sequencing of Herpes Simplex Virus 1 directly from human cerebrospinal fluid reveals selective constraints in neurotropic viruses

**DOI:** 10.1101/866350

**Authors:** Florent Lassalle, Mathew A Beale, Tehmina Bharucha, Charlotte A Williams, Rachel J Williams, Juliana Cudini, Richard Goldstein, Tanzina Haque, Daniel P Depledge, Judith Breuer

**Author notes:** These authors contributed equally (Florent Lassalle, Mathew A Beale). Department of Medicine, New York University School of Medicine, New York, New York, USA.

## Abstract

Herpes Simplex Virus type 1 (HSV-1) chronically infects over 70% of the global population. Clinical manifestations are largely restricted to recurrent epidermal vesicles. However, HSV-1 also leads to encephalitis, the infection of the brain parenchyma, with high associated rates of mortality and morbidity. In this study, we performed target enrichment followed by direct sequencing of HSV-1 genomes, using target enrichment methods on the cerebrospinal fluid (CSF) of clinical encephalitis patients and from skin swabs of epidermal vesicles on non-encephalopathic patients. Phylogenetic analysis revealed high inter-host diversity and little population structure. By contrast, samples from different lesions in the same patient clustered with similar patterns of allelic variants. Comparison of consensus genome sequences shows HSV-1 has been freely recombining, except for distinct islands of linkage disequilibrium (LD). This suggests functional constraints prevent recombination between certain genes, notably those encoding pairs of interacting proteins. Distinct LD patterns characterised subsets of viruses recovered from CSF and skin lesions, which may reflect different evolutionary constraints in different body compartments. Functions of genes under differential constraint related to immunity or tropism and provide new hypotheses on tissue-specific mechanisms of viral infection and latency.

## Introduction

Herpes simplex virus 1 (HSV-1, *human herpesvirus 1,* subfamily *Alphaherpesvirinae*) is a globally distributed human pathogen that chronically infects approximately 70% of adults and commonly causes mild skin vesicles (‘cold sores’) and mucosal urogenital ulcers. HSV-1 infection is characterised by lifelong latent infection of different parts of the nervous system, including trigeminal ganglia, ciliary ganglia (Lee et al. 2015), but also the brain (Yao et al. 2014), punctuated by periodic lytic reactivation and migration to the skin to form fresh vesicles (Higgins et al. 1993).

HSV-1 is a leading cause of viral encephalitis worldwide, with an estimated incidence of 250,000-500,000 cases per year, and mortality approaching 70% (Steiner & Benninger 2013; Hjalmarsson et al. 2007). An association of HSV encephalitis (HSE) with homozygous autosomal recessive mutations in interferon signalling pathways has been reported in young children (Gnann & Whitley 2017; Herman et al. 2012; Shen-Ying Zhang et al. 2013). However, in adults the majority of HSE occurs in individuals with intact adaptive immunity and no evidence of underlying deficiency of innate immunity. Encephalitis is also a feature of other neurotropic alphaherpesvirus infections including Equine Herpes Viruses (EHV), which causes respiratory disease, abortion and neurological disorders in horses. In this pathogenic system, viral genetics are demonstrably involved in encephalitis: a single nucleotide polymorphism (SNP) - the D752 variant of EHV1 viral polymerase (Nugent et al. 2006) – is associated with invasion and inflammation of the horse central nervous system (CNS), increased viraemia *in vivo*, and leukocyte tropism and reduced susceptibility to the anti-polymerase drug aphidicolin *in vitro* (Goodman et al. 2007).

Only a limited analysis of global HSV-1 sequence diversity has been performed to date (Szpara, Gatherer, et al. 2014; Kolb et al. 2013), notably because of difficulties in sequencing viruses from clinical material in a high-throughput manner. The viral genome size (152kbp) makes large-scale PCR amplicon-based sequencing studies extremely challenging while sequencing without enrichment yields insufficient data. Consequently, some HSV-1 genomes have been sequenced using PCR amplicons (Kolb et al. 2011; Szpara 2014; Watson et al. 2012), while other HSV-1 sequences were obtained from *in vitro* cultured isolates, followed by metagenomic sequencing (Szpara, Gatherer, et al. 2014; Szpara, Tafuri, et al. 2014; Kolb et al. 2011; Szpara et al. 2010). Cultured samples are associated with genetic bottlenecks causing a loss of intra-host diversity (Depledge et al. 2011). A more recent approach is the use of RNA/DNA-probe based hybridisation, or target enrichment, to sequence viral genomes directly from clinical specimens without the use of a culture step (Depledge et al. 2011; Houldcroft et al. 2017); this has been applied to a number of viral pathogens (Thomson et al. 2016; Palser et al. 2015; Depledge et al. 2014), including HSV-1 (Ebert et al. 2013; Greninger et al. 2018).

In this study we hypothesised that HSV-1 viruses causing HSE would demonstrate a genomic signal of neurotropism that delineates them from viruses causing ‘classical’ skin HSV-1. To examine whether polymorphisms in HSV-1 may be associated with increased neurotropism and encephalitis we recovered and sequenced HSV-1 genomes from the cerebrospinal fluid (CSF) of eight patients diagnosed with encephalitis and compared the results with HSV-1 genomes recovered from swabs of patients with acute cutaneous HSV-1 infections. We used targeted enrichment to enable direct sequencing of the viral population at the site of sampling (Houldcroft et al. 2017), and performed phylogenetic and population genomic analyses to describe the inter- and intra-host diversity in these patients. We detail the evidence of widespread recombination in HSV-1, the high homogeneity of population structure across samples from different vesicles on a same patient, and the distinct signatures of genetic linkage associated with strains derived from CSF and with strains derived from skin swabs.

## Methods

### Samples

All clinical samples were either obtained from patients treated at Royal Free Hospital (RFH; London) or submitted for reference analysis from regional laboratories to Public Health England (PHE; Manchester). Ethical approval for viral sequencing was obtained through the UCL Infection DNA Bank Fulham REC 12/LO/1089. The sample set represented 21 unlinked patients, comprising thirteen samples from cerebrospinal fluid (CSF) and fifteen from swabs or whole blood (SWAB) (Table 1). We note that no paired CSF and SWAB samples were available from a same patient. This is because on one hand sampling CSF is an invasive procedure only done in the extreme case of a declared encephalitis, and thus no CSF sample was collected on patients presenting skin lesions; on the other hand, none of the encephalitic patients presented skin lesions.

**Table 1:** Sample metadata and genome sequencing information.

| Patient | Patient Age (years) | Patient Sex | Sequence | Run Accession | Collection Year | Sample Type | qPCR Cycle Threshold (C<sub>t</sub>) | Sequencing Run | Total Reads | Mapped Reads (%) | Mean Coverage (all sites) | Genome coverage >250x (%) | Genome coverage >1x (%) | Used for Consensus Sequence Analyses | Used for Minor Variant Analysis |
| --- | --- | --- | --- | --- | --- | --- | --- | --- | --- | --- | --- | --- | --- | --- | --- |
| N1 | 39 | M | HSV1-SWAB1 | ERR3316613 | 2016 | Oral Swab | 33.2 | 2 | 1,727,324 | 74.4 | 1439.6 | 95 | 100 | Y | Y |
| N1 | 39 | M | HSV1-SWAB2 | ERR3316614 | 2016 | Whole Blood | 21.9 | 2 | 448,114 | 1.8 | 9.3 | 0 | 95.4 | N | N |
| N1 | 39 | M | HSV1-SWAB3 | ERR3316615 | 2016 | Genital Swab | 22.5 | 2 | 2,007,980 | 75.3 | 1705 | 96.1 | 100 | N | Y |
| N2 | 41 | F | HSV1-SWAB4 | ERR3316616 | 2016 | Facial Swab | 32 | 2 | 647,594 | 49.8 | 373.7 | 58.6 | 100 | Y | N |
| N3 | 15 | M | HSV1-SWAB5 | ERR3316617 | 2016 | Occular Swab | 25.6 | 3 | 2,662,700 | 74.8 | 2280.4 | 97.5 | 99.9 | Y | Y |
| N4 | 4 | M | HSV1-SWAB6 | ERR3316618 | 2016 | Skin Swab | 28 | 3 | 1,308,978 | 63.3 | 953.2 | 87.9 | 100 | Y | Y |
| N6 | 65 | F | HSV1-SWAB7 | ERR3316619 | 2016 | Genital Swab | 23 | 3 | 4,025,804 | 74.3 | 3354.1 | 98.7 | 99.8 | Y | Y |
| N7 | 0 (10 days) | F | HSV1-SWAB8 | ERR3316620 | 2016 | Oral Swab | 28.49 | 3 | 1,220,154 | 2.7 | 35.5 | 0 | 98.8 | N | N |
| N7 | 0 (10 days) | F | HSV1-SWAB10 | ERR3316622 | 2016 | Occular Swab | 23.2 | 3 | 6,362,766 | 79.9 | 5155.2 | 99.2 | 99.9 | Y | Y |
| N7 | 0 (10 days) | F | HSV1-SWAB9 | ERR3316621 | 2016 | Occular Swab | u/k | 3 | 3,265,926 | 71.2 | 2687.1 | 97.9 | 100 | N | replicate of HSV1-SWAB10 |
| N8 | 58 | M | HSV1-SWAB11 | ERR3316623 | 2016 | Oral Swab | 24 | 3 | 7,207,750 | 73.3 | 4994.9 | 99.3 | 99.9 | Y | Y |
| N8 | 58 | M | HSV1-SWAB12 | ERR3316624 | 2016 | Whole Blood | 31.7 | 3 | 1,344,718 | 1.7 | 24.2 | 0 | 98.6 | N | N |
| N9 | 38 | M | HSV1-SWAB13 | ERR3316625 | 2016 | Genital Swab | 29.7 | 3 | 1,469,206 | 46.2 | 792.4 | 85.8 | 99.9 | Y | Y |
| N10 | 15 | M | HSV1-SWAB14 | ERR3316626 | 2016 | Facial Swab | 24.3 | 3 | 2,910,694 | 73.5 | 2458.7 | 97.9 | 100 | Y | Y |
| N11 | 69 | M | HSV1-SWAB15 | ERR3316627 | 2016 | Oral Biopsy | 23.8 | 3 | 3,956,988 | 72.9 | 3278.5 | 97.9 | 99.9 | Y | Y |
| C1 | 57 | u/k | HSV1-CSF1 | ERR3316628 | 2013 | CSF | 26.8 | 1 | 2,296,834 | 77.8 | 2045.8 | 97.3 | 99.9 | Y | Y |
| C2 | 90 | u/k | HSV1-CSF2 | ERR3316629 | 2013 | CSF | 26.76 | 1 | 1,122,058 | 79.1 | 1016.6 | 94.3 | 99.9 | Y | Y |
| C3 | 61 | u/k | HSV1-CSF3 | ERR3316630 | 2013 | CSF | 30.04 | 1 | 1,094,248 | 47.0 | 594.3 | 87 | 99.7 | Y | Y |
| C4 | 80 | u/k | HSV1-CSF4 | ERR3316631 | 2013 | CSF | 30.19 | 1 | 1,940,454 | 63.1 | 1418.3 | 94.7 | 99.9 | Y | Y |
| C4 | 80 | u/k | HSV1-CSF7 | ERR3316632 | 2013 | CSF | 32.75 | 2 | 818,418 | 53.7 | 498.4 | 72.5 | 99.9 | N | replicate of HSV1-CSF4 |
| C4 | 80 | u/k | HSV1-CSF8 | ERR3316633 | 2013 | CSF | 30.39 | 2 | 828,236 | 57.8 | 544 | 75.8 | 100 | N | replicate of HSV1-CSF4 |
| C5 | 46 | u/k | HSV1-CSF5 | ERR3316634 | 2013 | CSF | 27.47 | 1 | 1,393,282 | 76.8 | 1224.2 | 94.3 | 99.9 | Y | Y |
| C5 | 46 | u/k | HSV1-CSF6 | ERR3316635 | 2013 | CSF | 31.35 | 2 | 1,226,662 | 55.3 | 752.2 | 81.1 | 99.8 | N | replicate of HSV1-CSF5 |
| C6 | u/k | u/k | HSV1-CSF9 | ERR3316636 | 2013 | CSF | 31.77 | 2 | 622,628 | 0.9 | 5.3 | 0 | 81.8 | N | N |
| C7 | 77 | u/k | HSV1-CSF10 | ERR3316637 | 2013 | CSF | 34.22 | 2 | 591,728 | 28.3 | 181.9 | 24.2 | 99.9 | Y | N |
| C8 | u/k | u/k | HSV1-CSF11 | ERR3316638 | 2013 | CSF | 34.95 | 2 | 570,486 | 0.8 | 6.5 | 0 | 88.3 | N | N |
| C9 | 66 | u/k | HSV1-CSF12 | ERR3316639 | 2012 | CSF | 33.03 | 2 | 584,314 | 29.6 | 190 | 26.8 | 99.9 | Y | N |
| C10 | 76 | u/k | HSV1-CSF13 | ERR3316640 | 2012 | CSF | 33.66 | 2 | 759,246 | 7.6 | 62.7 | 1.3 | 99.6 | Y | N |

u/k: unknown

Total DNA was extracted from 28 samples using QIAamp DNA Mini Kit (Qiagen) for CSF samples, QIAsymphony DSP virus/pathogen Kit (Qiagen) for blood samples, or NucliSENS EasyMAG (Biomerieux) for swabs or biopsy samples. Viral load was determined for CSF isolates by qPCR (Table 1). For patients N1, N7 and N8, samples from different sites were submitted for sequencing. For patients C4 three aliquots of the same sample (technical replicates) were analysed and for patients C5 and N7 biological replicates of the same site (CSF and ocular respectively) were analysed.

### Sequencing

Overlapping 120-mer RNA baits (2x tiling) were designed to selectively hybridise HSV-1 genomic DNA based on a multiple sequence alignment of 85 HSV-1 sequences and 9 HSV-2 sequences, representative of global sequence diversity obtained from GenBank as previously described (Depledge et al. 2011). Custom designed baits were uploaded to SureDesign and synthesised by Agilent Technologies.

DNA extracts containing HSV-1 DNA were quantified using the HS dsDNA assay for Qubit 3.0 (Thermo Fisher Scientific). Samples were prepared to obtain an input amount of 200-500ng or, where quantity was below 200ng, bulked to 200ng using human genomic DNA (Promega). All samples were sheared for 150 seconds using a Covaris E220 (Duty Factor 5%, Peak Incident Power 175 and 200 Cycles per Burst). Library preparation and hybridisation were performed according to the SureSelect^XT^ Target Enrichment System for Illumina Paired-End Multiplexed Sequencing Library protocol (Agilent) using a Bravo Automated Liquid Handling Platform Option B (Agilent). All libraries were sequenced in the same way, using 12 cycles of pre-capture PCR and 18 cycles of post-capture PCR, giving a total of 30 cycles. All recommended quality steps were performed. Final libraries were quantified by Qubit and normalised to equimolar concentrations.

Libraries were prepared in three batches as part of a routine automated workflow used for library preparation of multiple pathogens with organism-specific RNA baits (Depledge et al. 2011, 2014). Replicate samples were randomly distributed across three MiSeq runs, with some replicates together in the same run and others separated (see Table 1). Sequencing was performed on an Illumina MiSeq to generate 250-bp paired-end reads. Base calling and sample demultiplexing were generated using BaseSpace (Illumina).

### Genome Mapping and *de novo* Assemblies

The aim of this work was to compare variants between many genomes; to facilitate population-level analyses of genomes, we primarily opted for a reference mapping (rather than *de novo* assembly) approach – this enabled all variants to be related to one another reliably, but precluded analysis of repetitive regions or insertions/deletions. However, to ensure our analysis was not overly biased by reference mapping, we compared variant calls with the results of *de novo* assembly (as described below).

HSV-1 sequencing reads were extracted from the sequencing data by matching to a BLAST (Altschul et al. 1990) database, mapped to the HSV-1 strain 17 reference genome (NC_001806.2) (Davison 2011), and variants called at sites with >5 reads coverage using snippy (Seemann 2015) (v3.0), an automated pipeline that parallelises bwa (Li 2013), samtools (Li et al. 2009), and freebayes (v1.0.2) (Garrison & Marth 2012) for variant calling using GNU Parallel (Tange 2011). To call consensus SNPs, we required >50% of reads in agreement, and reads marked as having secondary mapping to more than one site were discarded before variant calling. We excluded problematic repeat genes (RL1, RL2, RS1) as these contain extensive short sequence repeats (SSRs) (Szpara, Gatherer, et al. 2014), which negatively impact upon accurate short-read alignment. This resulted in a final alignment of 138,278 nucleotide positions, of which 2,928 varied between sequences. We defined a subset of ‘core SNPs’ (variant positions covered by at least 5 independent sequencing reads in all genomes studied) using snippy-core (Seemann 2015), yielding a set of 2,734 well defined variant positions. We further refer to this alignment as the “mapped alignment” (available online at https://doi.org/10.6084/m9.figshare.10290470).

*De novo* assembly was initially performed on the HSV-1 reads using IVA (Hunt et al. 2015), and contigs were evaluated using the built-in quality control tools, with quast (Gurevich et al. 2013) and with Kraken (Wood & Salzberg 2014) to screen for contamination (including human genome). After evaluation, three IVA assemblies (HSV1-CSF10, HSV1-CSF12, HSV1-SWAB4) performed poorly, and we reassembled the reads using metaSPAdes (Nurk et al. 2017). To analyse our data in a global context, we aligned our *de novo* assembled contigs along with previously published sequences from around the world (Szpara, Gatherer, et al. 2014) to the HSV-1 strain 17 reference genome using parsnp (Treangen et al. 2014); this restricted the dataset to an alignment of 93,475 common, non-repetitive nucleotide positions including 2,135 parsimony informative sites. We further refer to this alignment as the “assembly core alignment” (available online at https://doi.org/10.6084/m9.figshare.8088605).

### Comparison of consensus genomes

For phylogenomic and recombination analysis of consensus genomes, we used only one sample per patient, and for patients with multiple samples, we selected the sample with the highest sequence coverage, making a dataset of 18 genomes (8 CSF, 10 SWAB) (Table 1). For phylogenomic analyses, this dataset was complemented with 25 publicly HSV-1 genome sequences (available at http://szparalab.psu.edu/hsv-diversity/data/) from a study of HSV-1 global diversity (Szpara, Gatherer, et al. 2014), for a final dataset of 43 genomes.

Based on the assembly core alignment, phylogenetic trees were produced with IQ-TREE (Nguyen et al. 2015) (v1.5.5) using a GTR gamma model optimised using the built-in model test, with 1,000 ultrafast parametric bootstraps and plotted using FigTree (Rambaut 2012) (v1.4.3). Neighbour Network analysis was performed using SplitsTree (Huson & Bryant 2006) (v4.14) with 1,000 bootstraps.

In the absence of strong population structure, we aimed to detect site alleles that would be associated with CSF or swab-derived (SWAB) samples. We converted our assembly core alignment into SNP-only variant files using snp-sites (Page et al. 2016) and vcf-tools (Danecek et al. 2011) v0.1.11, followed by analysis using PLINK (Purcell et al. 2007) v1.07 to test for SNPs association with either body compartment, applying the built-in adjustment for multiple testing.

To determine the extent of linkage disequilibrium (LD) acting on the clinical HSV-1 genome, we used our mapped alignment, which had a fuller coverage. Using LIAN (Haubold & Hudson 2000) v3.7, (available at http://guanine.evolbio.mpg.de/cgi-bin/lian/lian.cgi.pl/query), we estimated the genome-wide association between sites. We tested 2,735 polymorphic sites over 100,000 Markov Chain Monte Carlo (MCMC) iterations. LIAN outputs a summary statistic for standardised index of association (*I^S^_A_*) as a measure of the genome-wide linkage, with complete LD equalling zero, and a *p*-value indicating confidence.

To screen for potential localised genomic recombination, we performed a site-by-site LD analysis within 3000bp sliding windows of the same mapped alignment using the “genome-wide_LD_scan.r” script from the “genomescans” suite (Lassalle et al. 2016) (available at https://github.com/flass/genomescans). This scan tests all the association between all pairs of polymorphism patterns within a window using Fisher’s exact tests. Windows that contained less a minimum number (20) of biallelic sites were excluded; windows containing more than this number of SNPs were subsampled, selecting the most evenly spaced sites. To identify windows with stronger LD than in average genome-wide, it then compares the distribution of pairwise p-values within the window to the distribution obtained combining all possible windows in the genome using a Mann-Whitney-Wilcoxon test, reporting windows with - log_10_ p-value as a score, the local LD index (LDI, only scores > 5 were considered significant) (Lassalle et al. 2016). This was performed on an alignment of genomes from all patient-deduplicated samples in this study, as well as on sub-alignments of genomes covering the CSF and SWAB samples only, treating them as separate populations. To verify that the estimates of linkage in these data subsets reflected a group-specific population structure and not a bias induced by the difference in dataset sizes, we resampled the genomic datasets, drawing 30 pseudorandom combinations of genomes the size of each dataset, sampling equally from CSF and SWAB group. This way, we obtained a baseline distribution of local LD at each genome site under the hypothesis of no group-specific population structure; for group-specific subset analyses, only LDI scores falling out of the 95% confidence interval of this simulated distribution were deemed significant. To compare the strength of linkage estimated from the CSF and SWAB samples, we normalised the *r*^2^ values for CSF and SWAB datasets by dividing the *r*^2^ at each locus by the median *r*^2^ for the opposite dataset. We then computed the difference between normalised *r*^2^ values to identify significant windows with a substantial difference of LD strength between CSF and SWAB. Genomic windows (and the associated estimated values or *r*^2^ and LDI) were assigned to a gene according to the reference position of the centre point of the window.

It was previously shown that HSV-1 could recombine with members of the sister viral species HSV-2 (Wertheim et al. 2014; Koelle et al. 2017). As such long-distance recombination could bias our analyses, we searched for evidence of recombinant HSV-2 sequences in our genome assemblies. We ran a scanning blast search of 500-bp segments of the mapped assemblies of our genomes (the assemblies with most complete coverage) against the reference genome sequences HSV-1 strain 17 and HSV-2 strain HG52 (NCBI RefSeq accessions NC_001806.2 and NC_001798.2, respectively) (Davison 2011). None of our genomes has any segment of sequence that is more similar to the HSV-2 reference than it is to the HSV-1 reference, ruling out an introgression of HSV-2 in our samples (Supplementary data available online at https://doi.org/10.6084/m9.figshare.10247996).

### Within-host population analyses

For analysis of within-host polymorphism, we considered only high coverage genomes and included comparisons of duplicate samples from the same patients, resulting in a dataset of 19 genomes (8 CSF, 11 SWAB) (Table 1).

For minor variant calling, we remapped reads to the strain 17 reference (NCBI RefSeq accession NC_001806.2) using bwa mem (Li 2013), removed duplicate reads using MarkDuplicates from Picard Toolkit (Broad Institute) (v2.2.4) and repeated the analysis in freebayes, using a minimum frequency threshold of 2% and a minimum of 2 independent supporting reads (achieved by requiring ≥5 reads with support from both strands) – in practice, due to uneven coverage, this meant that positions lacking ≥250 read coverage (11 of 18 samples had >90% 250x coverage) could not reliably detect all potential within-host polymorphisms. To compare within-host polymorphism variant counts between samples, we considered down-sampling to a defined read depth per sample, but uneven genome-wide coverage meant we could not guarantee good coverage at all positions this way. Instead, we used samtools (Li et al. 2009) (v1.3.1) to extract depth of coverage from the filtered BAM files, and the VariantAnnotation (Obenchain et al. 2014) (v1.20.0) package in Bioconductor (Gentleman et al. 2004) (v3.3) to evaluate all polymorphic sites in the VCF files across the 14 highest coverage samples, only considering polymorphic sites where >250 high quality (non-duplicate) reads were available for all 14 samples. All data were plotted using ggplot2 package (Wickham 2009) in R (R Core Team 2014).

Intra-host nucleotide diversity (π) was determined using the mixed effect model-based approach described previously (Goldstein et al. 2018).

### Data and software availability

All sequencing reads were deposited at EBI-ENA under BioProject accession PRJEB32480. All consensus sequences, assemblies, and phylogenetic trees are available from the Figshare online data repository at https://figshare.com/projects/Comparative_genomics_of_HSV-1_from_cerebrospinal_fluid_of_clinical_encephalitis_patients/63368. Scripts used in these analyses, raw data and intermediary data are available from Github online data repositories at https://github.com/matbeale/HSV1_recombination_selective_constraints and https://github.com/flass/genomescans.

## Results

### Virus sequencing from clinical material

Target enriched sequencing of 28 clinical samples from 20 patients (Table 1) yielded an average of 1,900,000 paired-end reads/sample, of which an average of 1,200,000 paired-end reads were specific to HSV-1 (50% ± 29 SD on-target reads). Mean HSV-1 genome coverage/sample was 1,400x (range 5-5,155x), with poorest coverage found in the two whole blood samples and two of the CSF samples. Excluding the repeat regions (20.5% of total reference genome length) we achieved >95% genome coverage at 1x read depth for 26/28 samples. Using our predetermined criteria, we were able to generate 20 consensus sequences with 5x read depth at 99% of positions, and a further three samples had 5x read depth at 90-98% of the genome. The remaining five sequences were not of sufficient quality and were omitted from further analyses. Our final working dataset (after removal of the lowest coverage genomes) comprised 23 samples from 18 patients.

Phylogenetic network analysis showed short inner branches and very long terminal branch lengths leading to the tips, i.e. showing deep divergence between strain lineages (Figure 1). However, while unlinked samples clustered apart, differing by >350 SNPs at consensus level, the four groups of replicate samples obtained from the same patients (Table 1) clustered by patient with a maximum 1 SNP difference at the consensus level within each group (Supplementary Figure 1).

**Figure 1.**
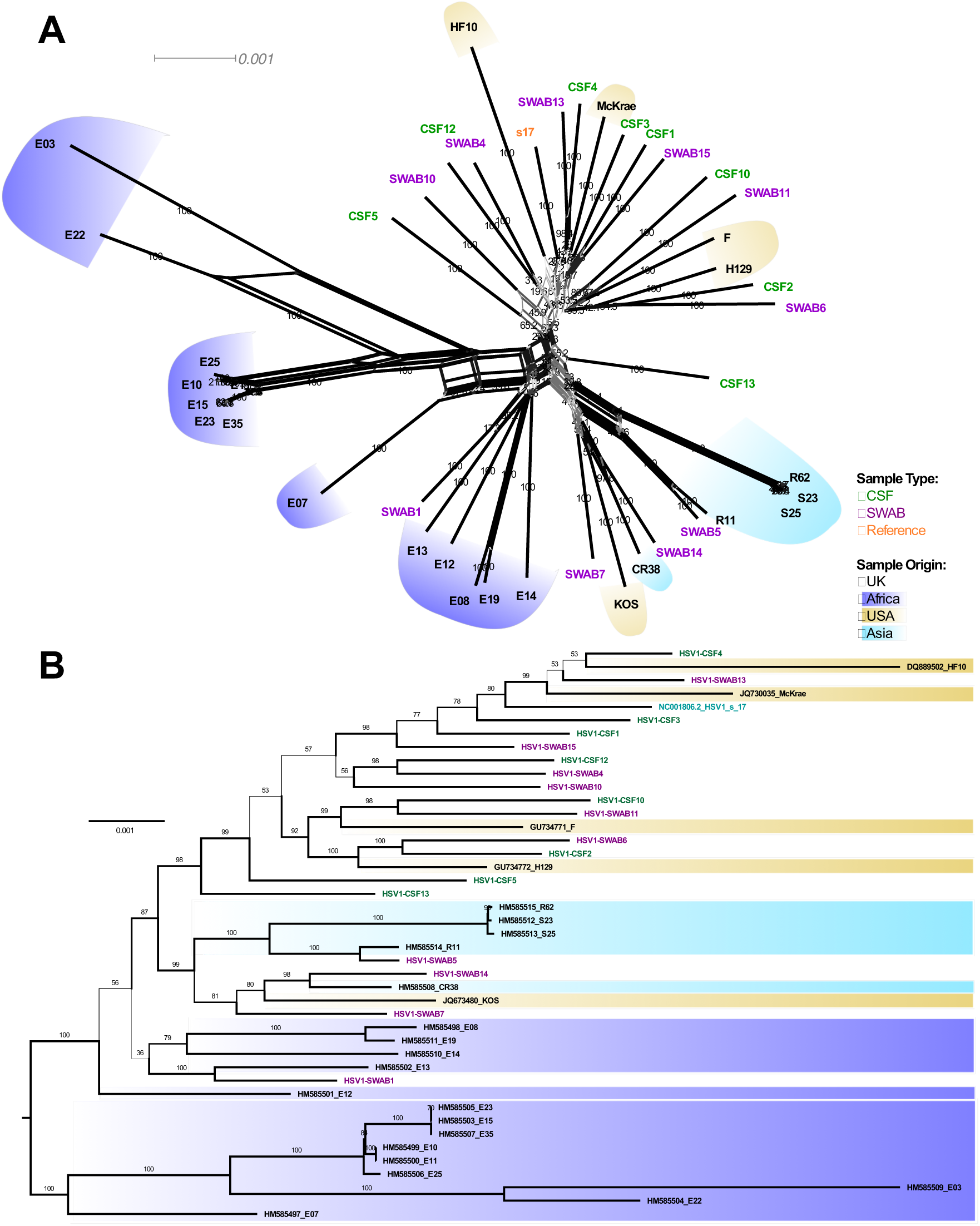
Phylogenetic analysis of unrelated HSV-1 genomes. Phylogenetic tree and network depicting the relationships of genomes obtained in this study and from Szpara et al. (2014). **A.** SplitsTree neighbour network (1000 bootstraps). Inner reticulation and ‘star-like’ branching pattern is indicative of extensive recombination. **B.** Maximum likelihood phylogeny (1000 ultrafast parametric bootstraps, midpoint rooted). Labels of sequences from this study are coloured by sample type, and background colour indicates geographical origin.

To further assure these data reflect the genuine viral diversity in samples, we compared the fine variation within samples. The frequency profiles of variants were thus compared between technical replicates of the same sample (i.e. a sample where multiple aliquots have been sequenced independently) from patient C4, as well as repeat samples from the same site i.e. biological replicates (N7 and C5) and two samples obtained from different sites in the same patient (N1; genital and oral). The number of unique variants (present in only one replicate) in the technical and biological replicates ranged from 2-15 (mean, 10; s.d., 4.7) which is likely to represent the error of the method (Supplementary Figure 2). Strikingly, the numbers of shared and unique variants in swabs from different sites (patient N1) was similar to that seen for technical and biological replicates. This suggests that the N1 genital and oral viral populations are closely related. We re-analysed variants with >1000x coverage in both samples from N1, to exclude potential errors associated with low coverage. The results maintained the original proportions of shared versus unique variants.

### Consensus-level genomic analyses

Of 138,278 nucleotide positions in the “assembly core alignment” of consensus genomes, we identified 2,928 variant sites. Roughly half (48%) of these sequence variants were singletons, i.e. limited to individual viral genomes. As described in previous studies (Szpara, Gatherer, et al. 2014; Brown 2004), a higher proportion of sites were polymorphic in the unique short (US) regions (3.0% sites) than the unique long (UL) regions (2.2% sites) (Supplementary Figure 4).

Since coverage may influence the ability to catalogue a variant (low coverage may lead a true variant to be discarded due to insufficient data), phylogenies generated from reference mapping approaches without coverage correction may be biased to appear closer to the reference. We therefore generated a ‘core SNP genome’ comprising only sites sequenced at the minimum coverage threshold of 5 reads in all samples. The resulting 2734 core SNPs were used as the basis of most of our downstream analyses.

### CSF and SWAB viruses cannot be discriminated based on phylogeny or unique variants

To contextualise our new genomes and to check whether CSF and SWAB genomes clustered as separate groups, our HSV-1 genomes were analysed together with other available HSV-1 genome sequences (Szpara, Gatherer, et al. 2014). The resulting phylogenetic network (Figure 1A) and phylogenetic tree (Figure 1B) both showed that most of our UK-sampled sequences clustered with other UK or US genomes. However, swab-derived samples from four patients (SWAB1, N1; SWAB5, N3; SWAB7, N6; SWAB14, N10) were phylogenetically distinct from our other UK sequences, clustering with a group of mostly Asian sequences. A principal component analysis (PCA) of variant positions considering our isolates only did not support significant population structure (Supplementary Figure 3). Considering clinical phenotype, phylogenetic analysis did not discriminate between CSF and SWAB samples (Figure 1A, 1B). This finding was confirmed by the comparison of multiple sequence alignments (having removed replicate patient samples) using PLINK, showing that no SNPs was significantly associated with CSF or SWAB group after p-value correction for multiple testing (data not shown).

### Evidence of pervasive recombination throughout the HSV-1 genome

Recombination has previously been reported for HSV-1 (Kolb et al. 2013; Norberg et al. 2011; Tang et al. 2014), and our SplitsTree analysis (Figure 1A) indicate significant genome-level recombination in this HSV-1 population (p < 0.01). To better characterise this pattern, we 1) used LIAN to describe the overall level of LD in our set of genomes, and 2) performed detailed scans of localised LD using a sliding window approach that identifies regions of the genome under increased LD. Note that these population genomic analyses were performed using only one sample per patient to avoid reporting structure from trivially clonal relationships in the dataset.

LIAN analysis of consensus-level variation in our HSV-1 dataset generated a standardised index of association (*I^S^_A_*) value of 0.0071 (P < 1×10^−5^), indicating very weak LD overall and suggesting that most loci are able to recombine freely. This contrasts even with other highly recombinant pathogens such as HIV-1, which has been shown to give an *I^S^_A_* of 0.0423 (Szmaragd & Balloux 2007), indicating that HSV-1 is more recombinogenic than HIV-1. In addition, we found no shared phylogenetic pattern in trees of distant genes (Supplementary Figure 5), confirming that, as previously described for other herpesviruses like human and murine cytomegaloviruses (Lassalle et al. 2016; Smith et al. 2013), the evolution of HSV-1 is characterised by pervasive recombination.

Given the very low linkage observed at the genome level, we next looked for potential local hotspots of LD, as potential indicators of evolutionary constraints. To detect loci which may be refractory to recombination, we applied genome-wide LD scans based on site-by-site test of association within short range (3,000 nucleotide windows), reporting the mean *r*^2^ value and assessing the significance of local peaks vs. the genome baseline with the local LD Index (LDI, see Methods). As for HCMV (Lassalle et al. 2016), hotspots of localized LD (here defined as the central position of genomic segments of 3,000 bp) occur in HSV-1 genomes (Figure 2). These LD hotspots, cluster predominantly in the unique short (US) region of the genome spanning US7/US8/US9/US10/US11/US12, as well as smaller and less intense hotspots of LD that cover genes UL9, UL47, and US3 (Figure 2A).

**Figure 2.**
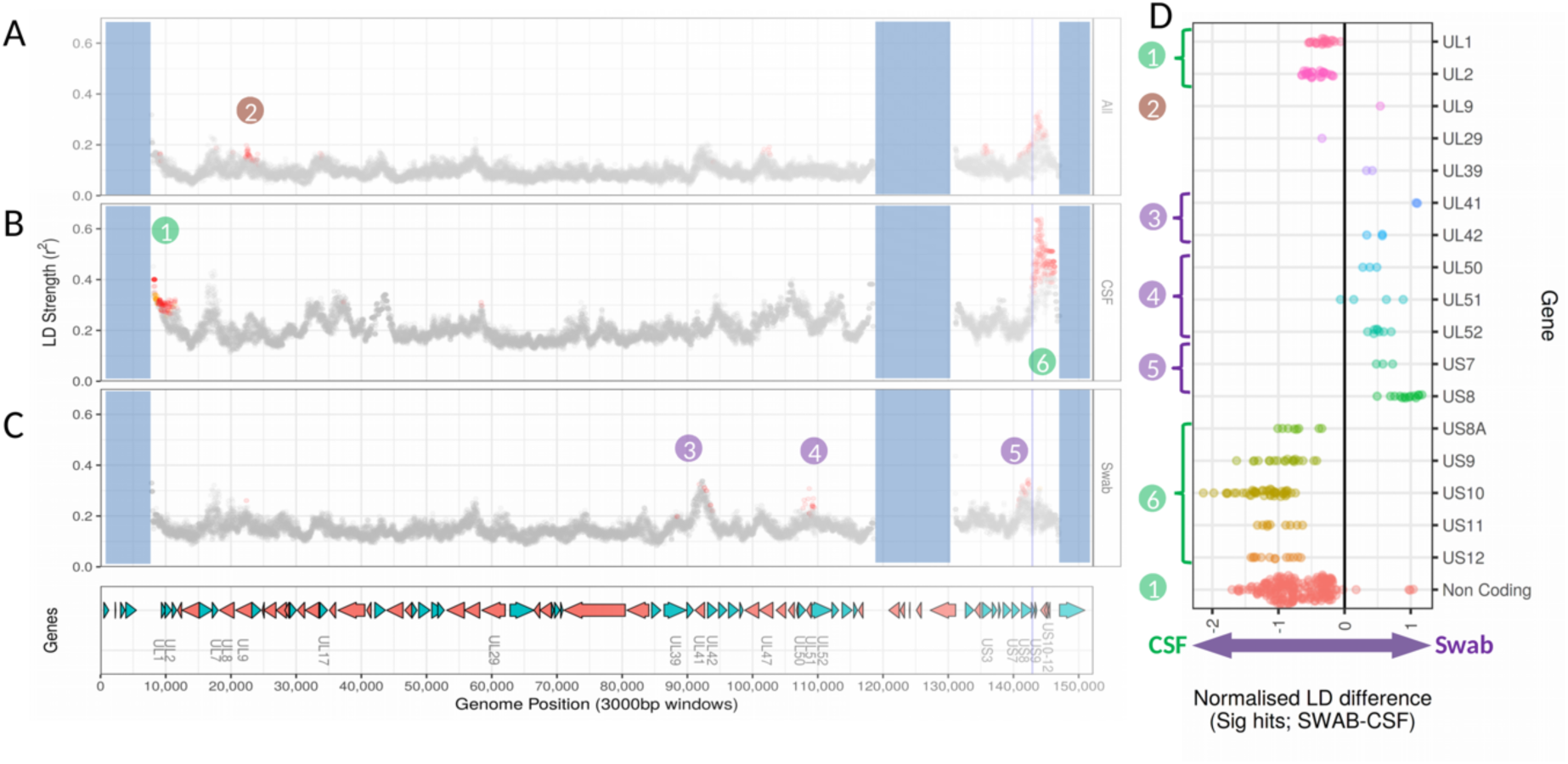
Variations in local intensity of linkage disequilibrium among consensus HSV-1 genome sequences derived from clinical samples. A sliding-window analysis of local linkage disequilibrium (LD) shows overall low linkage but reveals regions of increased LD, in particular in the US region. A-C plots show mean r^2^ (strength of local LD) measured in sliding windows (3000bp window span; incrementing by 10bp) along the reference genome, with significantly divergent sites (i.e. with local LD index > 5) highlighted in red. Vertical blue lines indicate start of US9 gene for clarity. Regions shaded in blue correspond to the repeated RL/RS genome regions (1-9338, 116926-132647, 145582-150962), excluded from the analysis. Analyses was performed for the entire dataset (A) and separately among genomes derived from CSF (B) and SWAB samples (C). For gene loci harbouring windows in significant local LD in at least one of these datasets, we computed the normalised difference of LD strength (D). Neighbouring genes with coherent signal are grouped in loci numbered 1 to 6, with colour indicating the dataset specificity of the signal of population structure: brown, whole dataset; green, CSF-specific; purple, SWAB-specific.

Considering CSF and SWAB sequences separately, we identified different patterns between the datasets (Figure 2B and 2C). LD analysis identified five potential genomic islands of local linkage where genes were inherited together in genomes derived from either CSF or SWAB samples, but not both (Figure 2D). The first islands of LD, namely genes UL1/UL2 and the upstream non-coding region (island 1, Figure 2B,D) together with the region covered by the genes US8A/US9/US10/US11/US12 (island 6, Figure 2B,D) had higher LD values in CSF samples. Other islands of LD covering genes UL41/42 (island 3, Figure 2C,D), UL50/51/52 (island 4, Figure 2C,D) and US7/US8 (island 5, figure 2C,D) had higher LD values in the swab-derived genomes. To test whether the differences of local LD intensities between the CSF and SWAB samples could have resulted from different genome sample sizes, we generated random genomic datasets containing the same number of genomes, but sampling equally from both groups (CSF and SWAB), and repeated the analyses over 30 permutations. At most positions in significant LD within CSF- or SWAB-only groups, the *r*^2^ value (LD intensity) for that group was also significantly above (out of the 95% confidence interval) the distribution drawn from these bootstrap permutations, showing that the five LD islands do indeed reflect specific population structure within the CSF and SWAB groups. Finally, a smaller peak of LD in gene UL9 was only detected when considering the whole dataset (island 2, Figure 2A,D), suggesting that population structure exists in this region irrespective of sample type.

We next examined the origin of these LD signals at the level of single SNPs, by analysing the site-to-site pattern of association underlying our scan (Supplementary Fig. 6; complete data available at Figshare; DOI: <u>10.6084/m9.figshare.8088635</u>). Because sites are often in local linkage with several of their neighbours (Supplementary Fig. 6D) – and in particular within islands of local LD – it is impossible to determine what SNP pair may be the underlying functional cause of the pattern of association. We therefore aimed to interpret the observed pattern in terms of interaction between proteins spanned by a same LD island (see Discussion).

### Within-host polymorphism frequencies are broadly similar between CSF and SWAB cohorts

Using only 364 sites with variant alleles with good coverage in all of the 14 samples (see Methods), we estimated the mean intra-host nucleotide diversity (π) for each sample, and recorded an average of 0.00027 (Figure 3B). Other than one outlier there was no difference in the number of minor variant sites per sample between the SWAB and CSF samples using the subset of variant positions (Figure 3A). The within-sample diversity for the outlier HSV1-SWAB7 (sequenced from a genital swab) was unexpectedly high, with 150 minor variant positions within our subset of robustly covered SNPs, and diversity was >5-fold greater than the mean of all other samples (*π*=0.0011 and *π*=0.0002, respectively). Genome-wide analysis of the within-sample polymorphisms showed that variants were distributed across the genome, with increased density between UL15-UL23, UL28-UL41 and in the US region (Figure 4). There was no clustering of variant frequencies around a discrete set of values, as could be expected in the case of multiple infection with distinct genotypes.

**Figure 3.**
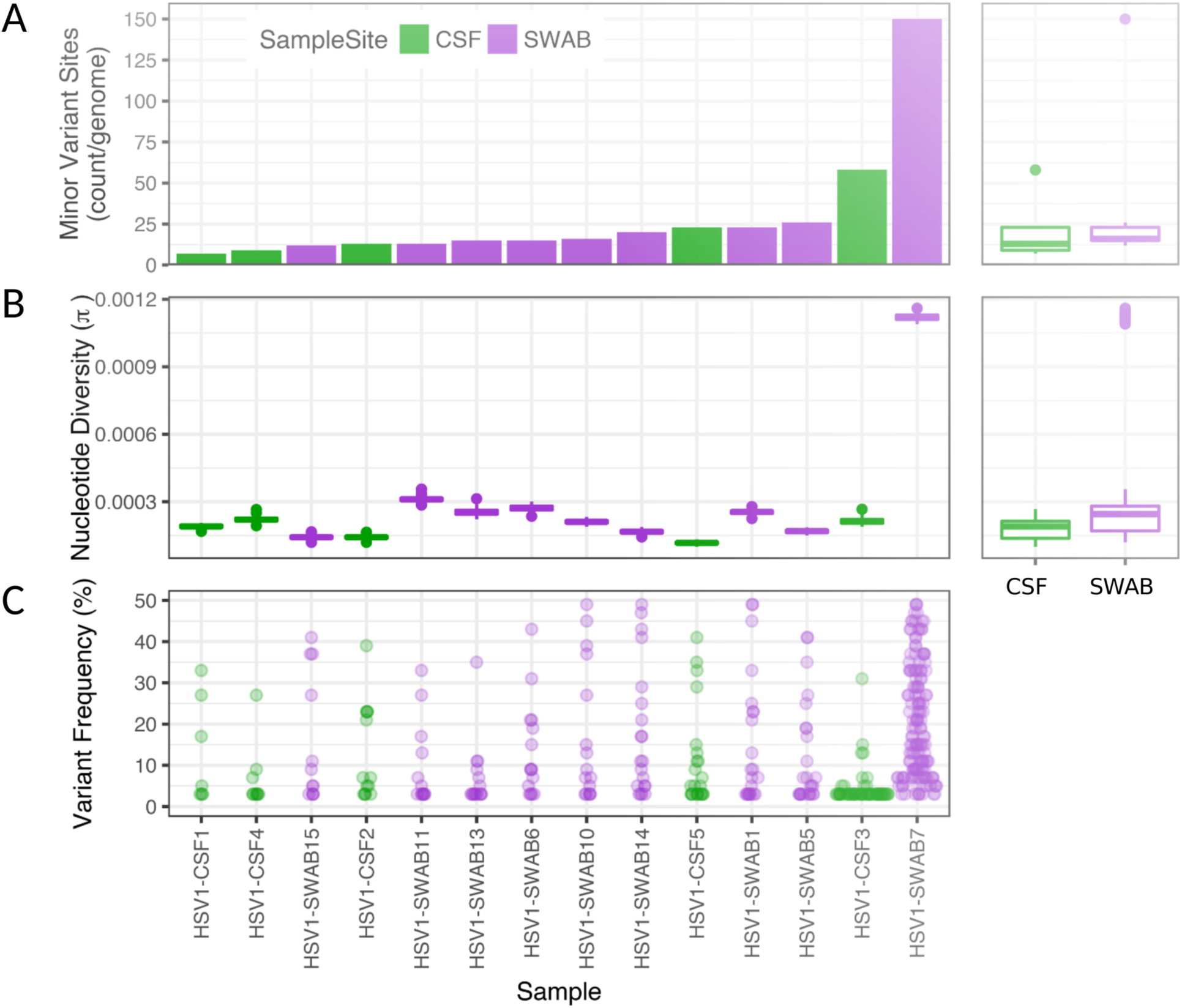
Within-sample analysis of variant genome positions. **A.** Density of sites with minor variants per sample, ordered from the last to the most heterogeneous sample. Most samples have <26 variant sites, with no difference in the relative number of variant sites between CSF and SWAB sample groups. **B.** Nucleotide diversity (π; estimated based on 100 re-sampled alignments) is low in all samples but HSV1-SWAB7, with no difference between CSF and SWAB samples. **C.** Most variants occur at low frequency, apart from in HSV1-SWAB7 where a large fraction of variants occur at intermediate frequencies.

**Figure 4.**
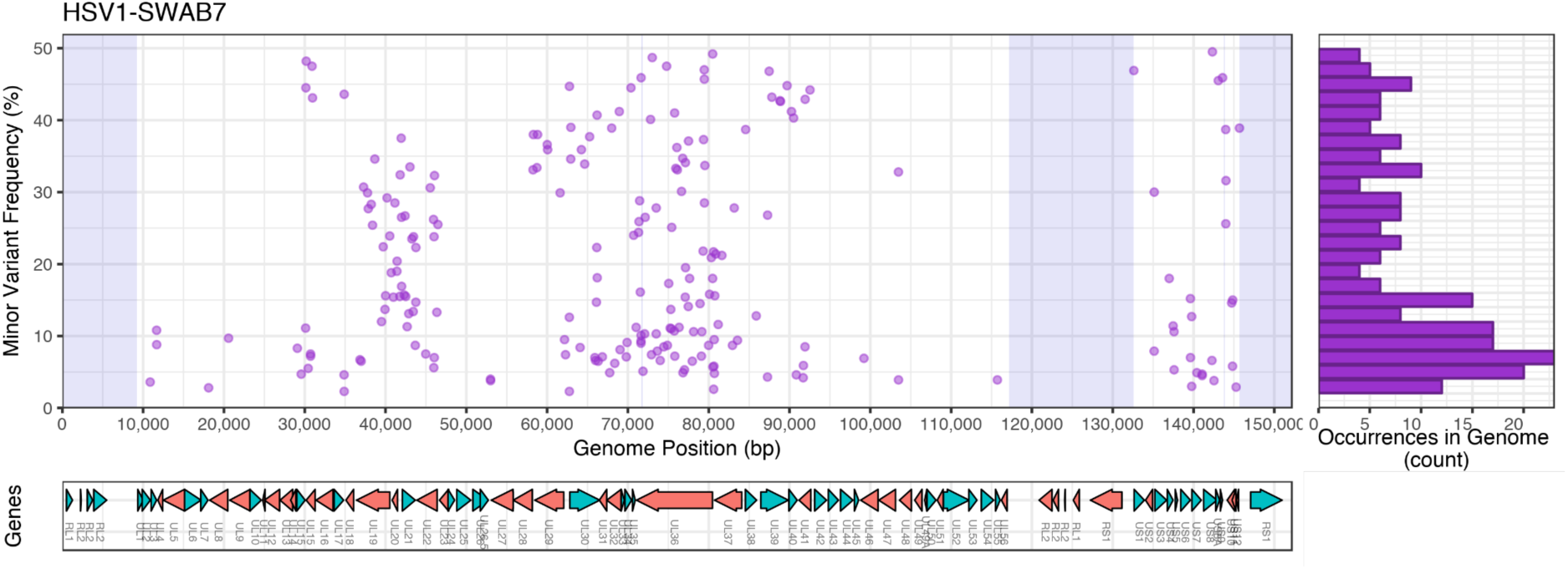
Frequency of HSV1-SWAB7 minor variants across the genome. Minor variant frequencies (estimated with a mean read depth of 3354x) show a mixed pattern of intra-host allelic diversity, with variants occurring at many different frequencies without clustering. This pattern suggests intense recombination between different virus types, possibly emerged from extensive intra-host evolution. Plots show frequency of variant alleles by genomic position (left) and frequency histogram (right, 2% intervals), with corresponding gene positions below. Shaded region indicates repeat regions with no mapped reads.

## Discussion

In this study, we aimed to investigate whether some HSV-1 types were more capable of causing encephalitis than others, as suggested by previous reports in equine herpes virus (Goodman et al. 2007). To test this hypothesis, we used target enrichment methods to sequence HSV-1 directly from patient samples without a culturing step, overcoming the associated problems of culture bias, population size reduction and loss of diversity (Houldcroft et al. 2017; Depledge et al. 2014; Greninger et al. 2018). This allowed us to recover for the first time high-quality draft HSV-1 genomes from encephalitis-derived samples. We also sequenced HSV-1 genomes from epidermal samples using the same methodology, so we could assemble a robust dataset generated under homogeneous conditions. We were able to obtain minimal whole genome coverage for 26/28 samples sequenced, and sufficient coverage for consensus genotype calling for 23/28 samples. Importantly, we had >250x coverage over most of the genome (sufficient for robust minor variant calling) in 14 samples. We show that this method is robust: blind resequencing of replicate samples taken from the same patient at the same time yielded consensus sequences varying by less than 2 SNPs, and those differences that were observed between consensus may be explained by variants which frequencies were close to 50% and flipped minor/major status between replicates. Within-sample population profiles were also broadly similar, with most polymorphic sites shared between replicate samples.

To understand the within-sample population variability of HSV-1 infection, we sequenced viral genomes in duplicate or replicate samples from four patients: two with encephalitis and two with skin vesicles (Figure 4). Although some variability was seen between technical and biological replicates, (Supplementary Fig. 2 C, D), it is likely explained by sequencing error, mapping ambiguity or insufficient coverage. Alternatively it is also possible that within-patient viral loads were sufficiently high so that our sampling was not fully representative of intra-host diversity (Illingworth et al. 2017) and suffered stochastic effects – a hypothesis that would be particularly likely for patient C5 where fixed differences between the biological replicates occur.

An interesting observation was the genetic homogeneity of HSV-1 at the variant level observed between samples obtained from swabs of different sites of the same patient (N1; Supplementary Figure 2 A, B). Previous modelling in VZV has indicated the existence of a bottleneck whereby individual vesicles are initiated by only 1-3 virions, whether during reactivation or primary infection and populations within each vesicle then evolve independently and develop a unique minority variant profile (Depledge et al. 2014). The high degree of similarity between the polymorphism profile of populations from oral and genital sites in patient N1 is not consistent with such a scenario of population transmission with bottleneck and diversification. Firstly, the rarity of unique variants occurring between the populations sampled at both sites indicate they are related and had little time to diverge. There is however no means to infer here the nature and direction of their relationships: one may be parent of the other or both may be derived from another unsampled population, etc. Secondly, the close similarity of frequencies of shared variants suggests that the propagation of populations involved large enough numbers of virions so that the new-founded population was completely representative of its source. This pattern is consistent with the hypothesis of recent transfer of the viral population from one site to the other by auto-inoculation.

Among our samples from patients with skin lesions, the sample HSV1-SWAB7 had surprisingly high level of intra-host diversity; there were no clearly distinct haplotypes occurring in the sample, which would have resulted in the clustering of variant frequency around a discrete set of values – a phenomenon that could be observed in case of recent super-infection with separate viral lineages without significant gene flow (Cudini et al. 2019). The data we observe suggests another scenario - that different lineages were in contact within the host and that recombination occurred frequently between them, and then within their offspring population – enough to decorrelate the frequency of variants at different sites. The intermediate frequencies of variants imply they are still segregating in the population and the pattern we observed is likely a snapshot of a dynamic process. However, in the absence of longitudinal data, the time scale and directionality of this dynamic process, and whether it is close to an equilibrium, cannot be determined. Another – non-exclusive – possibility is that the viral population carries mutations in the DNA replication genes that confer a hyper-mutator phenotype, which would lead arising of numerous independent variants and their fast accumulation in the population. We found no truncating mutations in genes that are considered essential to the replication pathway (Weller & Coen 2012). However, non-synonymous substitutions occur in these replication genes, including four private to the hyper-diverse SWAB7 sample; some of the mutations occurring in UL30 appear not to be fixed in the sample (Sup. Table 1). These substitutions could have an impact on replication fidelity, but genetic investigations would be required to assess their functional role with respect to such phenotype.

As described in previous studies (Szpara, Gatherer, et al. 2014; Norberg et al. 2011), we found a lack of strong population structure (between unrelated samples), leading to star-like phylogenies with long terminal branches. Given an estimated HSV-1 global seroprevalence of >70% (meaning several billion persistently infected individuals), this sample of 18 UK patients represents a tiny fraction of global diversity, and one can only speculate at the level of recombination and ancestral mixing occurring in the unsampled majority. HSV-1 has very low linkage disequilibrium – far lower than most ‘classically’ recombining organisms (Szmaragd & Balloux 2007) and, as previously shown for human cytomegalovirus (Lassalle et al. 2016), most sites in the genome are essentially freely recombining. However, this contrasts with elevated linkage disequilibrium in specific genomic islands, particularly around the US region – a region where two out of nine genes are associated with early stage infection (Depledge et al. 2019). The fact that these genes are inherited as a haplotype more frequently than the rest of the genome suggests that they are prevented from recombining, possibly due to selective constraints.

We speculate that the selective pressures acting on viruses during CNS infection and encephalitis are different from those selective pressures met by viruses in canonical epithelial infection. In that case, a genomic signature of adaptation may exist in the pathogen population that is successful in invading the CNS niche and/or proliferating during encephalitis. This phenomenon has previously been observed for mumps vaccine virus isolated from a patient with encephalitis (Morfopoulou et al. 2017). In this dataset, HSV-1 genomes from CSF and SWAB samples did not segregate by phylogeny or by any specific variants. However, we found that recombination affecting a region spanning US7/US8 was under greater constraint in SWAB genomes. The products of US7 and US8 (gE, gI) are glycoproteins that together form a heterodimer, and have been shown to promote transport of HSV-1 glycoproteins towards the neuronal axon termini for viral assembly prior to spread into epithelial cells (Snyder et al. 2008; Rajcáni et al. 1994). In contrast, the US9 gene, which product has the related function of promoting transport of HSV-1 capsid proteins towards the neuronal axon termini (Snyder et al. 2008), is physically located next to the US8 gene but in our dataset did not appear to be in LD with US7/US8. The presence of strong linkage disequilibrium between two genes whose protein products directly interact might be expected, since conformational perturbation of the proteins mediated by genomic recombination could potentially impact on heterodimer formation. Since LD was specifically increased for the US7/8 region in SWAB genomes but not CSF genomes, it is possible that viruses that successfully transit from the neuronal axon to form epithelial vesicles have been selected for ‘pairs’ of gE/gI proteins, but that this selective constraint is relaxed when spreading in the CNS environment.

Within the SWAB patient group, we also found increased LD for a region spanning UL41/42 genes (LD island 3) and the UL50/51/52 genes (LD island 4) (Figure 2C,D and Supplementary Fig. 7). Genes in island 4, respectively encode a deoxyuridine 5’-triphosphate nucleotidohydrolase, a tegument protein and the viral DNA primase; the potential mechanism constraining their co-evolution remains unclear. Within LD island 3, UL42 encodes part of the DNA polymerase processivity factor complex and may also act as an inhibitor of innate immunity; UL42 has been shown to inhibit TNF-*α*-induced NF-*κ*B activation (Jie Zhang et al. 2013, 65). UL41 encodes the virion host shutoff (Vhs) protein, an endonuclease that cleaves host and viral messenger RNAs (mRNAs) during the lytic phase. This activity decreases host protein production, increases viral mRNA turnover and therefore facilitates the sequential expression of viral genes; it also impedes antigen presentation by the major histocompatibility complex (MHC) and inhibits the secretion of cytokines for the recruitment of peripheral adaptive and innate immune cells. Vhs has been previously reported to be associated with ocular and CNS infection in a mouse model (Brandt et al. 2003; Kolb et al. 2016). The key disease-associated mutation Kolb et al. reported at position 384 (N384S) is present in our dataset and the frequency of the disease-associated allele C (translating to amino-acid S) is higher in CSF samples then in SWAB samples (0.75 and 0.5, respectively), but the difference is not statistically significant (Supplementary Fig. 7). Importantly, it has been reported that the effect of UL41 on virulence was dependent on its association with genes UL42 and UL36/37, indicating an epistatic relationship between UL41 and its neighbour UL42 (Brandt et al. 2003). Vhs targets specific sites near the translation start codon, with an increased cleavage rate for mRNAs with optimal translation initiation sequence pattern (Shiflett & Read 2013). This sequence specificity, likely dependent on the Vhs protein structure, could also explain the co-evolutionary constraint between UL41 and the nearest transcribed gene UL42, which has an essential role in viral replication and immune evasion. While previous report suggests epistasis between UL41/42 is linked to virulence, we see stronger linkage in the SWAB group than in the CSF group (Supplementary Fig. 7). These observations are not contradictory, as the latency/epidermal reactivation/host transmission cycle on one hand and the neuro-invasion pathway on the other hand may both impose constraints on the co-evolution of UL41 and UL42, while selecting for different allelic combinations. Constraints driving stronger linkage in the SWAB group are potentially immune-driven, and may indeed be relaxed in the CNS, due to the different and smaller range of immune cells represented in this segregated compartment. However, the LD strength of LD island 3 peaks in UL41 (Figure 2C), suggesting constraints within Vhs protein may be the main driver of this local drop in recombination rate.

Within CSF samples, we found a region of large excess of LD spanning the non-coding region next to the RL repeat region and the genes UL1/UL2. UL1 encodes the envelope glycoprotein L (gL) that forms a complex with gH, which is essential for membrane fusion and cell entry. UL2 encodes an uracil DNA glycosylase that associates with the viral replisome. Here again there is no evidence linking the function of these genes; one possibility is that strong constraints against recombination within gH-encoding UL1 only accounts for the whole signal in this region, a hypothesis compatible with the quicker drop in LD outside the UL1 gene when shorter scanning window intensities are used (Sup. Mat. Online; doi: 10.6084/m9.figshare.8088635). Alternatively, because these gene share the same transcriptional unit (Depledge et al. 2019), which in addition can be fused with the transcript of RL2 located upstream in RL region, it might be postulated that the co-evolutionary constraint emerges from a – yet unidentified – transcriptional-level process.

We found the highest signal of LD in a region spanning US8A/9/10/11/12 of CSF genomes (island 6 in Fig. 2B,D). These genes are strongly associated together in CSF patients, suggesting a high level of evolutionary constraint, possibly caused by physical interaction of their protein products that is required for fitness in the CNS. US9 and US10 are tegument proteins involved in capsid synthesis and generally associated with the late phase of viral cellular infection and, as mentioned earlier, US9 also promotes anterograde capsid transport (Snyder et al. 2008). US11 binds DNA and RNA, interacting with US12/13/14 (Attrill et al. 2002). US12 encodes protein ICP47, which has a key role in host immune suppression via binding transporters associated with antigen processing (TAP), preventing MHC-based recognition of infected cells by cytotoxic T-lymphocytes (York et al. 1994; Früh et al. 1995). The specific activity of these immune cells in the brain tissue (McGavern et al. 2002) might create particular constraints for US12 function.

The US8A gene which is found at the edge of linkage islands 5 and 6 (Figure 2), is in high linkage with island 5 in SWAB samples and conversely with island 6 in CSF samples. Our LD scan uses 3-kb wide windows and its resolution is thus of the order of three genes, so it is possible that the LD signal in windows centred on US8A might have been driven by stronger LD in the neighbouring genes in each sample group. Intriguingly, US8A has been noted to be a potential virulence factor for neuro-invasion: US8A null mutants resulted in 10-fold increase in LD50 during CNS infection of mice (York et al. 1994; McGavern et al. 2002). Such a role in CNS infections might lead to strong structural constraints that could explain its association with the US9/10/11/12 gene block (island 6) in CSF samples.

In summary, we confirm that HSV-1 is highly recombinant and we show that, like HCMV, there are few regions of linkage disequilibrium with the genome. We find differences in the pattern of LD between HSV-1 sampled from peripheral sites as compared with CSF. In particular, strong compartment-related differences in the LD between genes in the US region are apparent and these effects are strong enough to be observed even when the data are considered as a whole (Figure 2A vs. Figure 2B,C). In addition, the phylogenetic relatedness between genomes of CSF and SWAB groups rules out the possibility that ancestral differences would account for the difference of population structure observed within each compartment. Although we do not have paired peripheral and CNS samples, we speculate that these distinct linkage constraints are likely to have arisen within the host. Given the limited size of our sample and the fact that all CSF patients were treated in the same (British) hospital, we cannot conclude on the generality of our findings. They may nonetheless represent some of the differential outcomes of selection elicited by the contrasting environment of epidermal vs. CNS body compartments. Whether the genomic regions under specific constraint in CSF genomes determine the ability of viruses to migrate to the CNS, or represent viral adaptation to the CNS environment once it has been colonized, warrants further investigation.

## Funding Information

This work was conducted as part of the PATHSEEK consortium (University College London, Erasmus MC, QIAGEN AAR, and Oxford Gene Technology), which received funding from the European Union’s Seventh Programme for research, technological development and demonstration (FP7) under grant agreement no 304865. We acknowledge joint Centre funding from the UK Medical Research Council and Department for International Development to the MRC Centre for Global

Infectious Disease Analysis (grant MR/R015600/1). FL was supported by the UK Medical Research Council (grant MR/N010760/1). MAB was supported by Wellcome funding to the Sanger Institute (098051). DPD was supported by a Medical Research Foundation New Investigator Award. JB receives funding from the NIHR UCL/UCLH Biomedical Research Centre. The funders had no role in the study design or decision to submit.

## Acknowledgements

The authors would like to thank the Royal Free Hospital Virology Department (London) and Public Health England (Manchester) for collection and extraction of samples, in particular Prof Ray Borrow for his help with ethics, and the staff of the UCL Pathogen Genomics Unit for performing the sequencing. The authors acknowledge the use of the following computational facilities in the completion of this work: the UCL Legion High Performance Computing Facility (Legion@UCL) and associated support services, the UCL Computer Science Department computer cluster and associated support services, and the Medical Research Council funded CLIMB project (www.climb.ac.uk) (Connor et al. 2016).

## Data Availability

Genomic data has been deposited to EBI-ENA under Bioproject accession PRJEB32480 and Run accessions ERR3316613–ERR3316640. Genome assemblies, multiple sequence alignments and phylogenetic trees are hosted on the Figshare online data repository at https://figshare.com/projects/Comparative_genomics_of_HSV-1_from_cerebrospinal_fluid_of_clinical_encephalitis_patients/63368; individual data items can be accessed with the following DOIs: doi: 10.6084/m9.figshare.8088605, doi: 10.6084/m9.figshare.8088635, doi: 10.6084/m9.figshare.8201813, doi: 10.6084/m9.figshare.8201828 doi: 10.6084/m9.figshare.8231093, doi: 10.6084/m9.figshare.10247996, doi: 10.6084/m9.figshare.10290470, doi: 10.6084/m9.figshare.10299314. Code used for analyses, raw data and intermediary data are available on Github at https://github.com/matbeale/HSV1_recombination_selective_constraints.

## Ethical statement

Ethical approval for viral sequencing was obtained from through the UCL Infection DNA Bank Fulham REC 12/LO/1089.

## Conflicts of Interest

The authors have no conflicts to declare.

## Abbreviations

HSV-1: Herpes Simplex Virus type 1
HSE: HSV encephalitis
CSF: cerebrospinal fluid
US: unique short
LD: linkage disequilibrium
LDI: local LD index

---

**Supplementary Table 1.** List of non-synonymous mutations observed in the consensus sequence of sample HSV1-SWAB7 with respect to the reference strain 17 genome sequence (NCBI RefSeq accession NC_001806.2).

| Gene | Genome pos. | CDS pos. | Prot. Pos. | Ref codon | Ref. AA | sample codon | sample AA | unique among CSF/SWAB samples? | reference allele occurs as minority var. at freq. |
| --- | --- | --- | --- | --- | --- | --- | --- | --- | --- |
| UL8 | 19420 | 1057 | 353 | GCC | A | GTC | V |  |  |
| UL8 | 18644 | 1833 | 612 | TGC | C | GGC | G |  |  |
| UL9 | NA |  |  |  |  |  |  |  |  |
| UL30 | 62903 | 96 | 33 | AGC | S | GGC | G |  |  |
| UL30 | 64080 | 1273 | 425 | AAC | N | ACC | T | unique | 0.084 |
| UL30 | 66101 | 3294 | 1099 | GCC | A | ACC | T |  | 0.147 |
| UL30 | 66144 | 3337 | 1113 | TCC | S | TGC | C |  | 0.223 |
| UL30 | 66177 | 3370 | 1124 | CCT | P | CAT | H |  | 0.407 |
| UL30 | 66197 | 3390 | 1131 | GCG | A | TCG | S | unique | 0.181 |
| UL30 | 66428 | 3621 | 1208 | ACC | T | GCC | A |  |  |
| UL5 | 14933 | 199 | 67 | CAT | H | CGT | R |  |  |
| UL5 | 14519 | 613 | 205 | TTG | L | TCG | S |  |  |
| UL5 | 14033 | 1099 | 367 | GAG | E | GCG | A | unique |  |
| UL42 | 94241 | 1129 | 377 | TTG | L | TCG | S |  |  |
| UL52 | 109680 | 631 | 211 | GTG | V | GCG | A |  |  |
| UL52 | 110589 | 1540 | 514 | GGC | G | GAC | D |  |  |
| UL52 | 110591 | 1542 | 515 | CCC | P | ACC | T |  |  |
| UL52 | 110735 | 1686 | 563 | CGC | R | TGC | C | unique |  |
| UL52 | 111221 | 2172 | 725 | GAT | D | AAT | N | unique |  |
| UL52 | 111282 | 2233 | 745 | GAA | E | GCA | A | unique |  |
| UL29 | 61925 | 129 | 44 | TCC | S | GCC | A |  |  |
| UL29 | 61012 | 1042 | 348 | TTC | F | TGC | C |  |  |

**Supplementary Figure 1.**
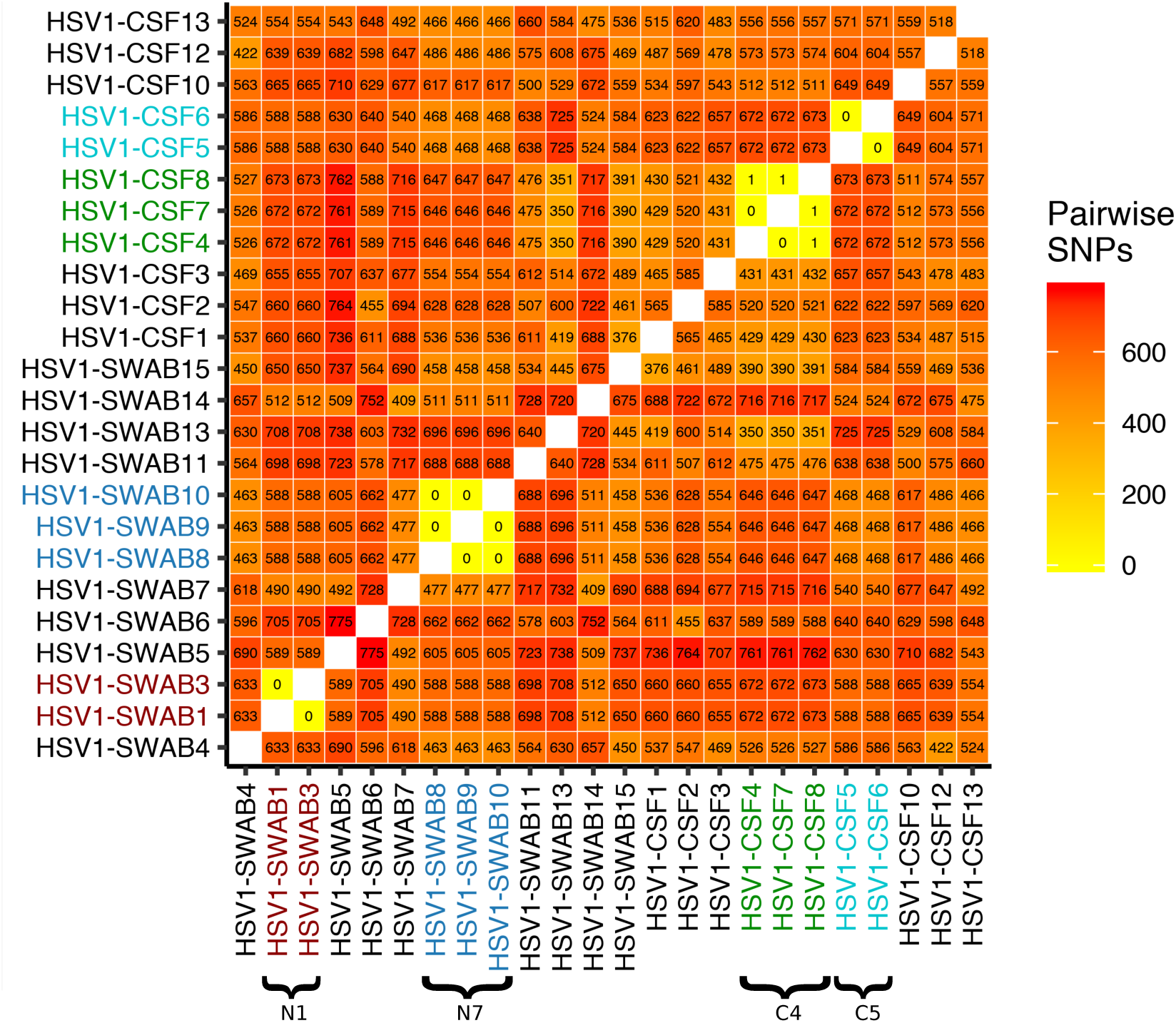
Pairwise SNP differences between unrelated samples and samples taken from the same patient. Unrelated samples are separated by >350 SNPs, while related samples have virtually identical consensus sequence (≤1 SNP) within the same patient. Samples from the same patient are indicated with brackets.

**Supplementary Figure 2.**
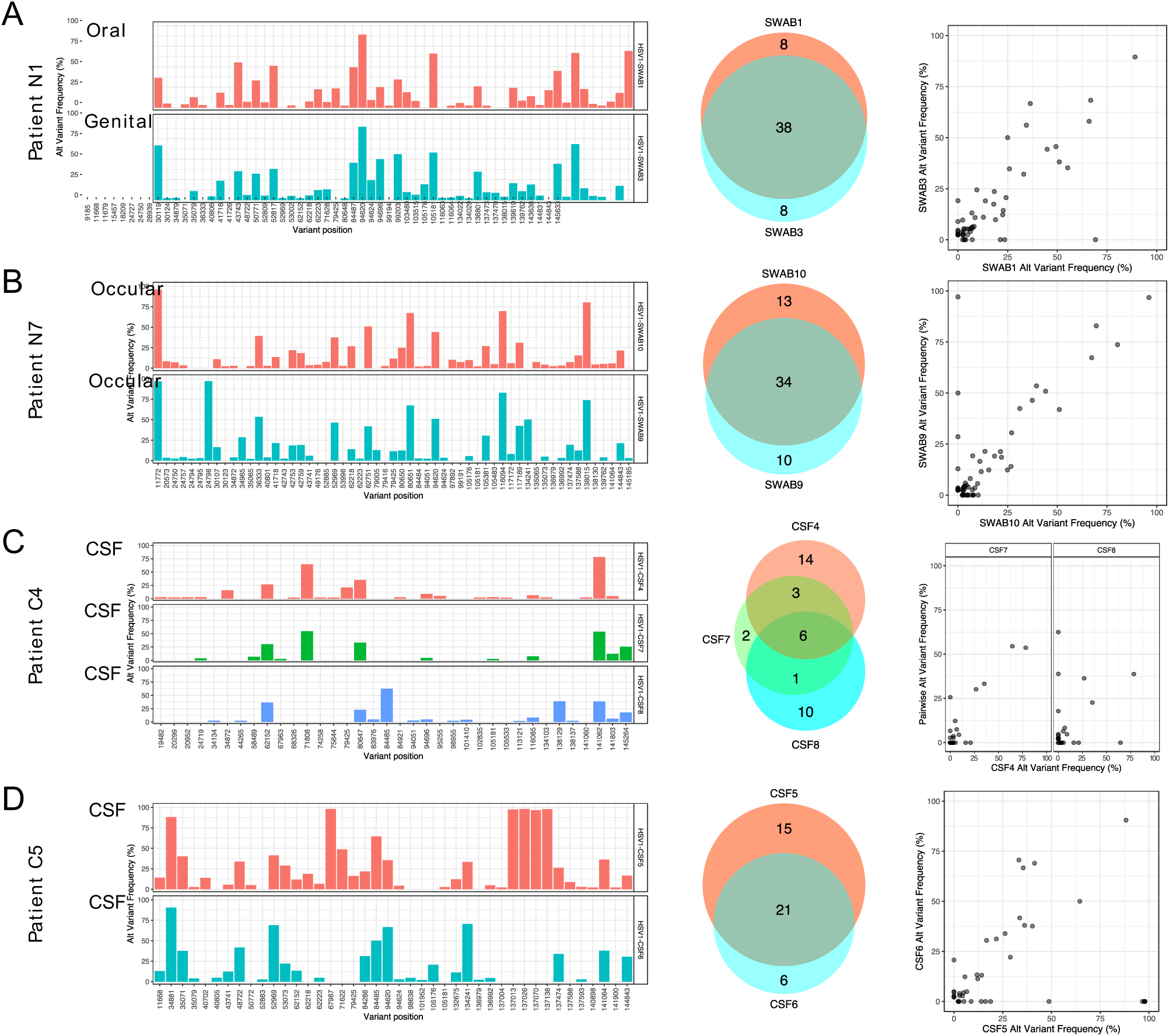
Replication of minor variant allele profiles between paired samples from same patient. Barplots show alternate (non-reference) allele frequencies by position, Venn diagrams indicate common minor variant positions, and scatter plots show frequencies of common variants in pairwise comparison. A. Patient N1, multiple sites. B. Patient N7, biological replicates from the same site. C. patient C4, technical replicates (same sample). D. Patient C5, biological replicates from the same site.

**Supplementary Fig 3.**
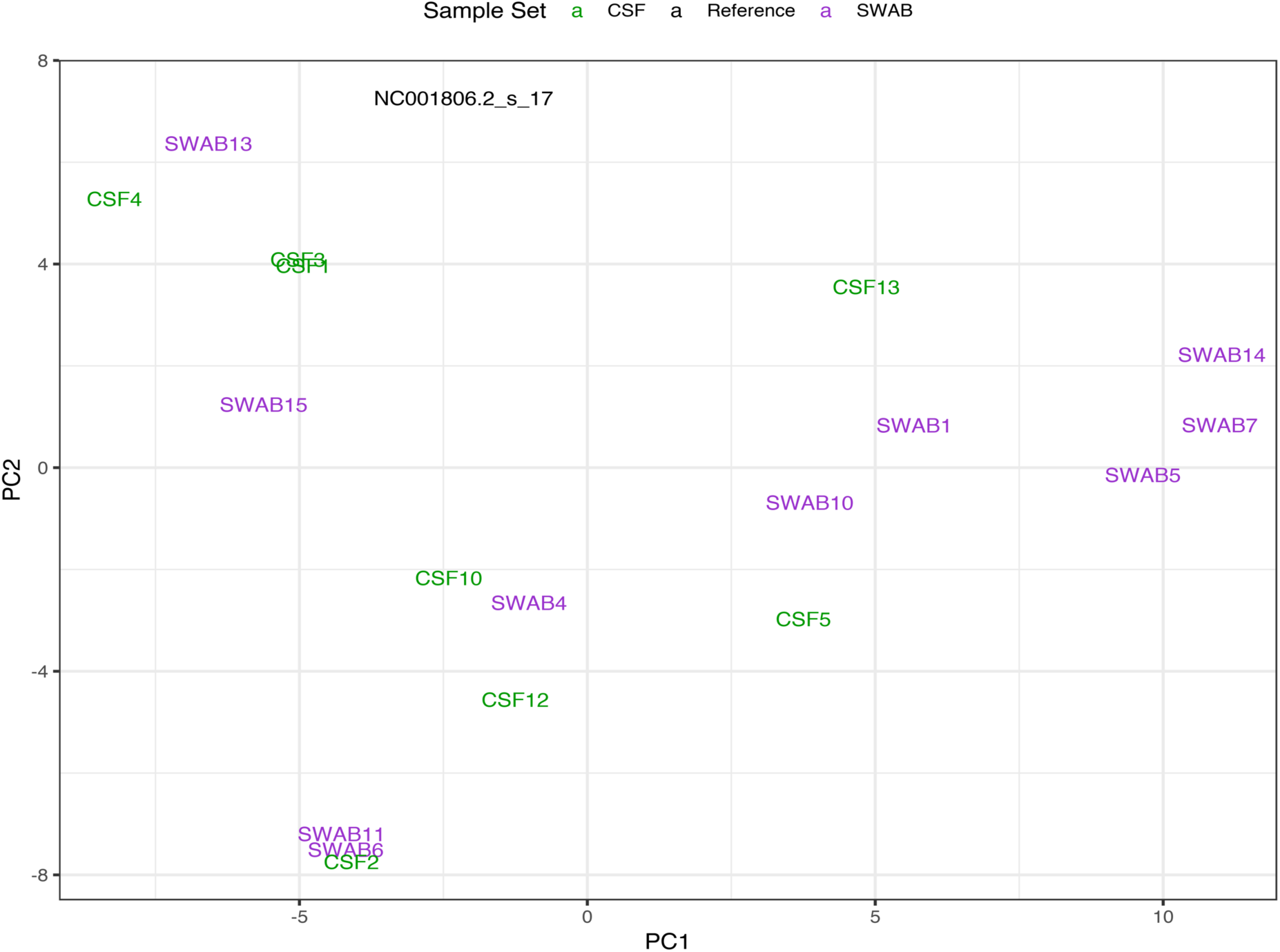
PCA of covariance of polymorphisms between consensus sequences shows absence of strong population structure. Samples coloured by sample category: green: CSF; purple: SWAB; black: reference.

**Supplementary Figure 4.**
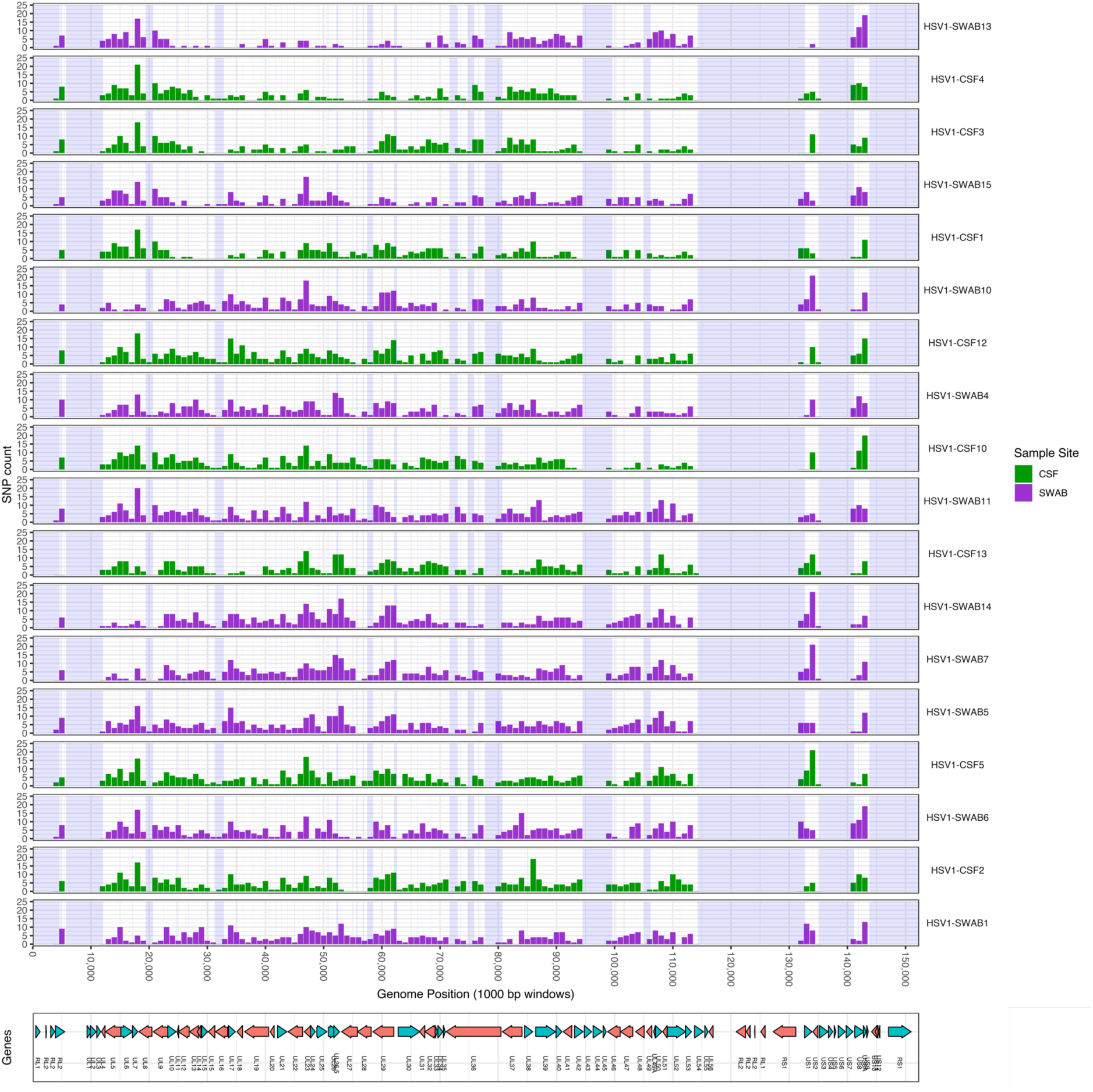
Genome-wide SNP density per sample. SNP density was plotted in sliding windows (500bp) along the genome in all samples from this study using the assembly core alignment. SNPs are broadly distributed along the genome, but increased diversity was observed in the US region. Gaps in the core-genome alignment are indicated with blue shade.

**Supplementary Figure 5.**
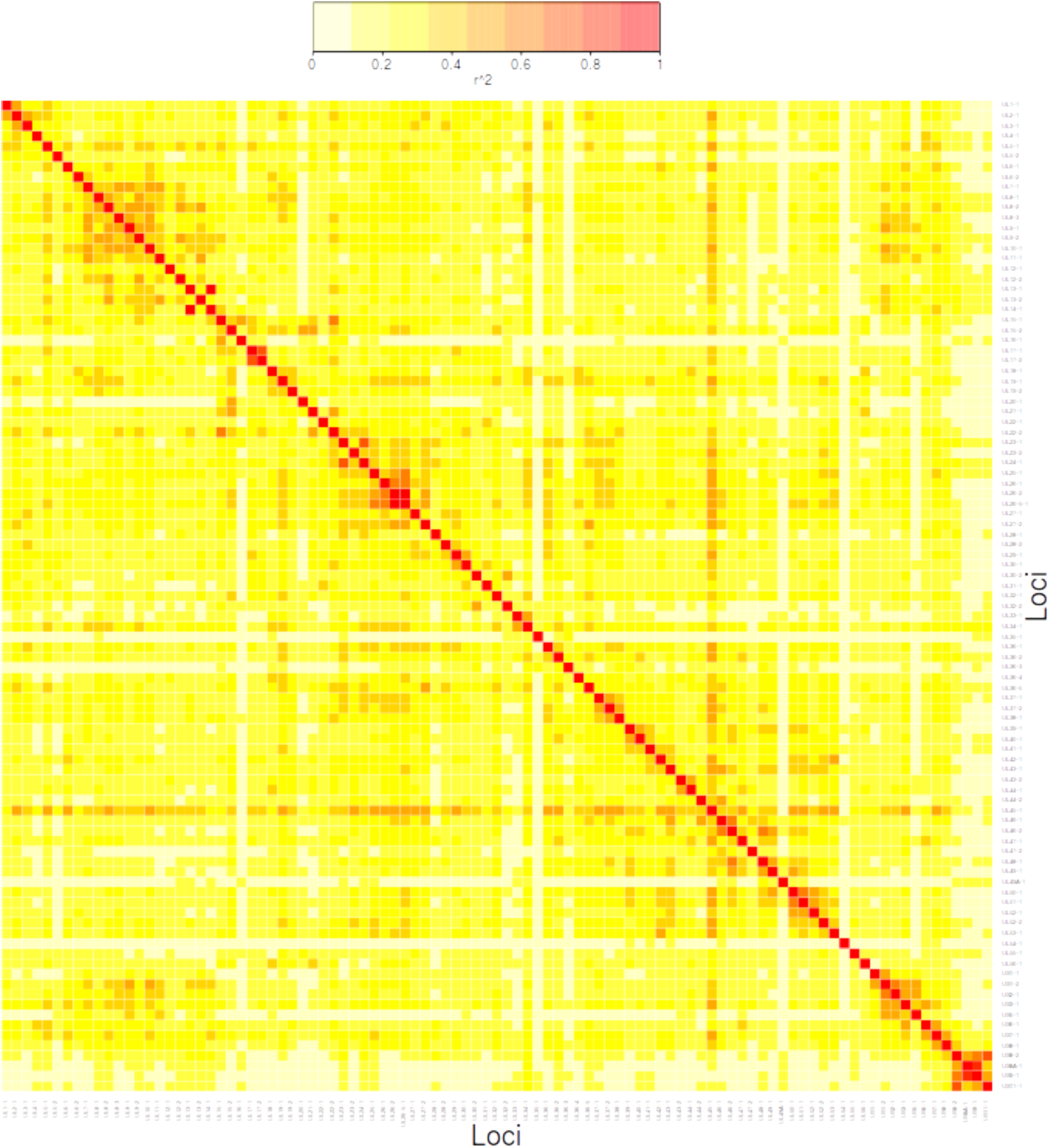
Gene-to-gene correlation of phylogenetic profiles of population structure. The phylogenetic relationships between consensus genomes were dissected for each gene locus using the bayesbipartprofile tool (https://github.com/flass/genomescans). Phylogenetic compatibility of tree splits with each gene alignment was computed for all splits observed in at least one Bayesian gene tree sample at a minimum frequency of 0.1 and separating clades of at least 3 sequences. The gene-to-split compatibility scores were then used to compute correlation (r^2^) between gene pairs, displayed as a heatmap; genes are ordered following their position in the genome. Genes physically clustered together show significant correlation of phylogenetic structure (i.e. related history of descent), but not those located far apart in the genome. Elevated compatibility of UL45 with most other genes indicates limited phylogenetic information available in this gene’s alignment.

**Supplementary Figure 6.**
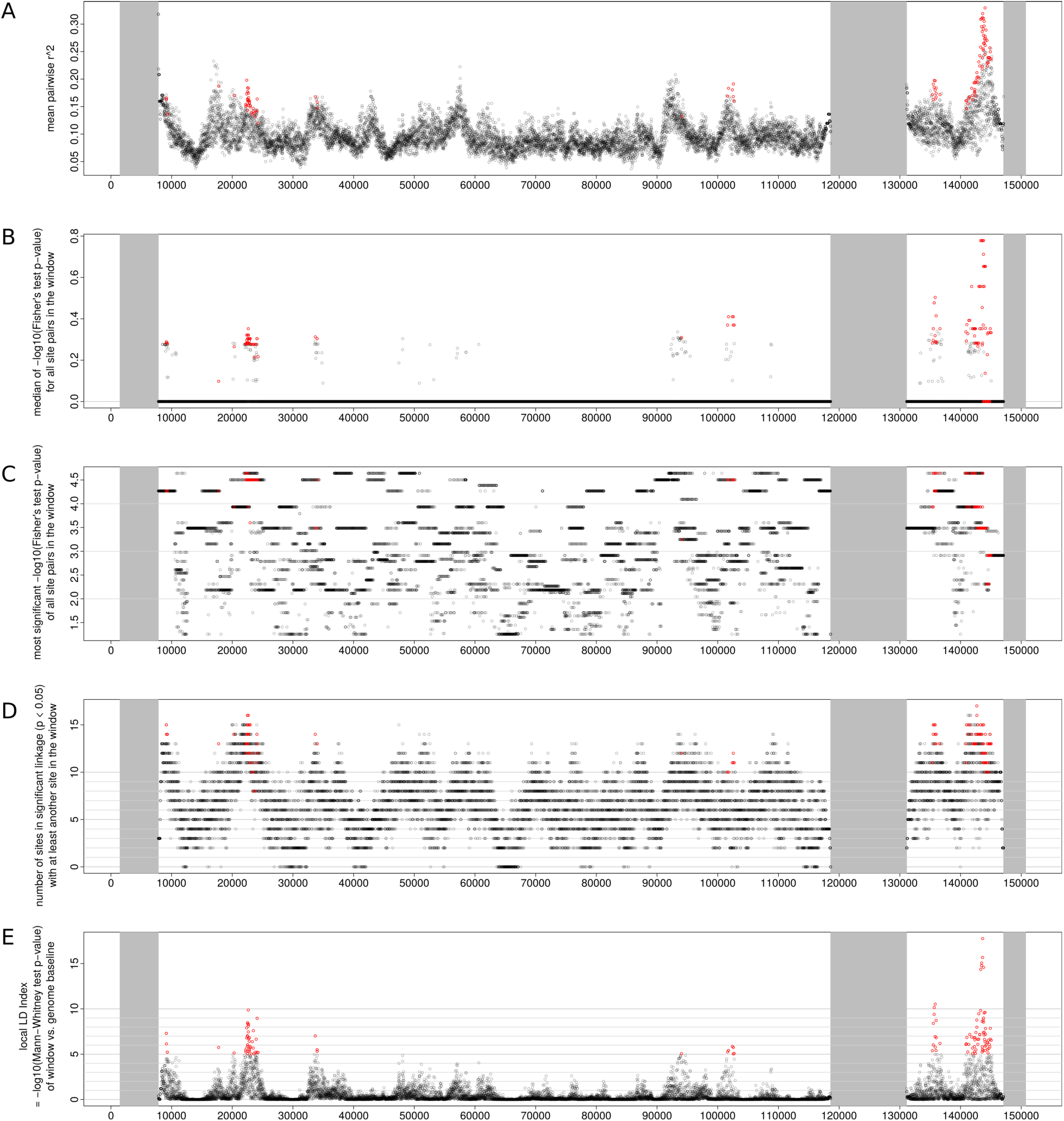
Comparison of local LD metrics in genome-wide scan of 18 CSF and SWAB HSV-1 genomes. A sliding window scan was conducted over the mapped alignment of 18 HSV-1 consensus genomes (3000bp windows, 10bp step; SNPs were subsampled so each window contains exactly 20 bi-allelic SNPs), and various metrics of within-window LD were reported, each aggregating in a different simpler LD metrics (*r*^2^ or Fisher’s exact test p-value) measured for each of the 190 pairs site present in a 20-site window): A) mean *r*^2^ as in Figure 2A; B) median Fisher p-value (-log_10_ transformed); C) most significant Fisher p-value (-log_10_ transformed); D) number of sites in significant linkage (Fisher p-value < 0.05) with at least another site within the window; E) local LD index (LDI), which is the - log_10_ transformed p-value of a one-sided Mann-Whitney-Wilcoxon test comparing the set of Fisher test p-values within the window to the distribution of values observed in all windows across the genome. Dots in red indicate the windows with LDI > 5, reported as significant local LD peaks in Figure 2A.

**Supplementary Figure 7.**
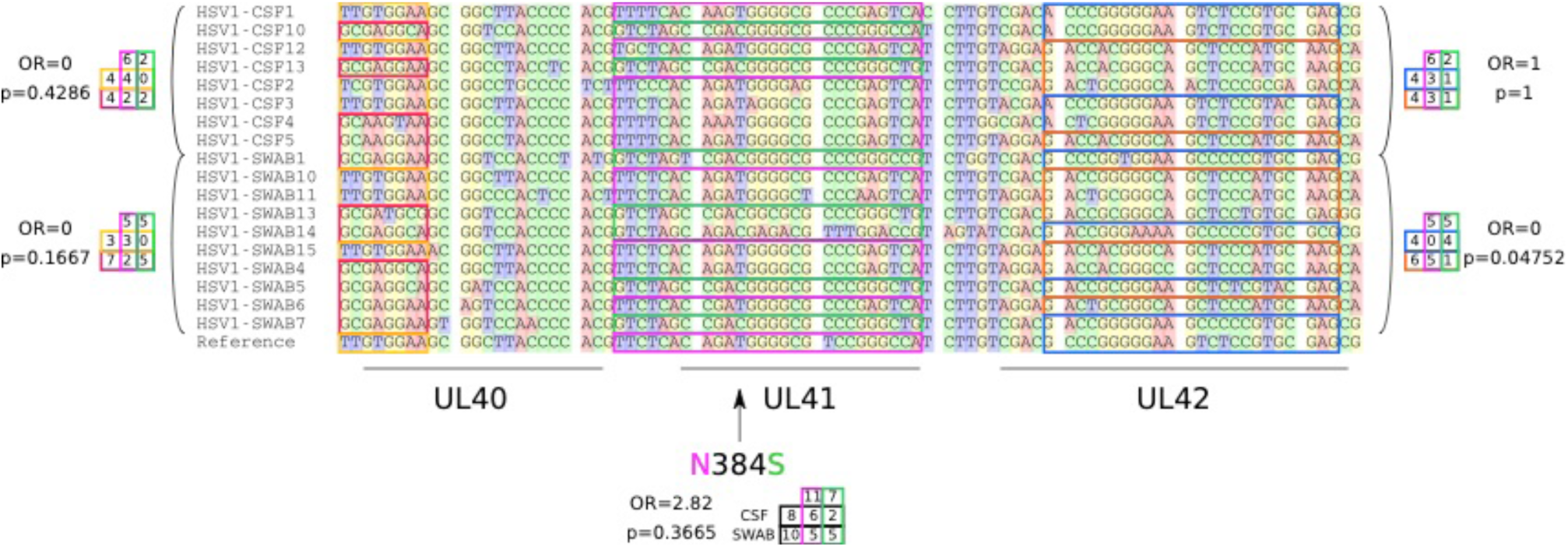
Haplotype structure of UL40/41/42 locus. Mapped alignment of consensus genomes from eight CSF and ten SWAB samples, reduced to the bi-allelic nucleotide sites occurring within the coordinates 89,900 to 94,600 (relative to reference strain 17), i.e. the region spanning the genes UL40, UL41 and UL42. Haplotypes, i.e. stretches of sites that are phylogenetically congruent, are framed and colour-coded according to a binary grouping per gene locus. Sites without framing were deemed subject to too much recombination to be classified into haplotypes. Agreement of haplotype structures between gene loci is indicated separately for the CSF (top) and SWAB (bottom) groups for UL40 vs. UL41 (left) and UL41 vs. UL42 (right), respectively. Results of Fisher’s exact tests for each group/comparison show that population structure is stronger across these loci within the SWAB group than within the CSF group. The non-synonymous site (genome position 91,486) leading to an asparagine to serine change (N384S) in UL41 protein product Vhs is indicated as it was previously reported to be associated with an increased virulence in a mouse model of ocular infection (Kolb et al. 2016).

## Notes

https://github.com/matbeale/HSV1_recombination_selective_constraints

https://figshare.com/projects/Comparative_genomics_of_HSV-1_from_cerebrospinal_fluid_of_clinical_encephalitis_patients/63368

